# Muscle specific translational control of Cand2 by mTORC1 regulates adverse cardiac remodeling

**DOI:** 10.1101/2020.11.29.403196

**Authors:** Agnieszka A. Gorska, Clara Sandmann, Eva Riechert, Christoph Hofmann, Ellen Malovrh, Eshita Varma, Vivien Kmietczyk, Lonny Jürgensen, Verena Kamuf-Schenk, Claudia Stroh, Jennifer Furkel, Matthias H. Konstandin, Carsten Sticht, Etienne Boileau, Christoph Dieterich, Hugo A. Katus, Shirin Doroudgar, Mirko Völkers

**Affiliations:** Department of Cardiology, Angiology and Pneumology, University Hospital Heidelberg, Im Neuenheimer Feld 410, 69120 Heidelberg, Germany; DZHK (German Centre for Cardiovascular Research), partner site Heidelberg/Mannheim, Germany; Medical Research Center, Medical Faculty Mannheim, Heidelberg University, Mannheim, Germany; Section of Bioinformatics and Systems Cardiology, Department of Cardiology, Angiology, and Pneumology and Klaus Tschira Institute for Integrative Computational Cardiology, University of Heidelberg, Heidelberg, Germany

**Keywords:** Cand2, Cardiac, Hypertrophy, mTOR

## Abstract

The mechanistic target of rapamycin (mTOR) is a key regulator of pathological remodeling in the heart by activating ribosomal biogenesis and mRNA translation. Inhibition of mTOR in cardiomyocytes is protective, however, a detailed role of mTOR in translational regulation of specific mRNA networks in the diseased heart is largely unknown. A cardiomyocyte genome-wide sequencing approach was used to define mTOR-dependent post-transcriptional gene expression control at the level of mRNA translation. This approach identified the muscle-specific protein Cullin-associated NEDD8-dissociated protein 2 (Cand2) as a translationally upregulated gene, dependent on the activity of mTOR. Deletion of Cand2 protects the myocardium against pathological remodeling. Mechanistically, we found that Cand2 links mTOR signaling to pathological cell growth by increasing Grk5 protein expression. Our data suggest that cell-type-specific targeting of mTOR might have therapeutic value for adverse pathological cardiac remodeling.

## Introduction

Pathological cellular remodeling is a hallmark of heart failure (HF) independent from the underlying etiology such as pressure overload, myocardial infarction, or inherited cardiomyopathies. Several studies and our recent work revealed that pathological cardiac stress induces early morphological changes and cardiac hypertrophy due to the activation of the kinase mechanistic target of rapamycin complex 1 (mTORC1)(Buss *et al*, 2009; Völkers *et al*, 2013; Sciarretta *et al*, 2018; Zhang *et al*, 2010). mTORC1 promotes protein synthesis by induction of rRNA transcription, ribosomal protein synthesis, phosphorylation of translation initiation factors and enhances translation of specific mRNAs that are required for stress adaptation (Ma & Blenis, 2009),(Iadevaia *et al*, 2012). Mechanistically, mTORC1 promotes translation of a specific subset of mRNAs regulated by the translation initiation factor eIF4E by direct phosphorylation of the eIF4E binding proteins (4E-BPs) (Thoreen *et al*, 2012),(Hsieh *et al*, 2012). mTORC1-sensitive transcripts often contain a terminal oligo-pyrimidine (TOP) or TOP-like motif in the 5’ UTR and mTORC1-dependent transcripts are predominantly regulated on the translational level (Jefferies *et al*, 1994) (Avni *et al*, 1996),(Thoreen *et al*, 2012).

Systemic pharmacological or genetic mTORC1 inhibition prevents pathological hypertrophy and improves cardiac function in murine disease models (Buss *et al*, 2009; Shioi *et al*, 2003; Völkers *et al*, 2013), but no established therapeutic regime targets mTORC1 at the level of cardiomyocytes in patients yet. Moreover, the role of mTORC1 in translational regulation of specific mRNA networks in the diseased heart is largely unknown, partly because existing tools in previous studies were unable to analyze gene expression at the level of translation.

We aimed to characterize mTORC1-dependent changes in gene expression that mediate cardiac response to pathological stress. Ribosome profiling (Ribo-seq) was used to obtain quantitative measurements of translation, and to dissect the mTORC1-dependent translational regulation of gene expression in cardiomyocytes. Among other genes dependent on the activity of mTORC1 was the uncharacterized muscle-specific gene Cullin-associated NEDD8-dissociated protein 2 (Cand2). Initially named transcription factor TATA-binding protein 120B (TIP120B), Cand2 belongs to the TIP120 gene family and shares 60% homology with TIP120A (Cand1) (Aoki *et al*, 1999).

In contrast to ubiquitously expressed Cand1, Cand2 is a muscle-specific protein and has been identified only in mammals. Both proteins have been shown to directly bind and modulate the activity status of Cullin1 (Cul1) (Liu *et al*, 2002),(Goldenberg *et al*, 2004),(Shiraishi *et al*, 2007). Cul1 is a component of SCF-like E3 ligase complex that promotes ubiquitination (Zheng *et al*, 2002),(Furukawa *et al*, 2000),(Zou *et al*, 2018). The activation of the SCF complex is stimulated by reversible and covalent post-translational modification (neddylation) of CUL1, which relies on ligation of the ubiquitin-like polypeptide Nedd8 protein to a specific lysine residue in the C-terminus (Goldenberg *et al*, 2004),(Min *et al*, 2005).

We found that pressure overload induced Cand2 expression on the level of translation which depends on the activity of mTORC1. Cand2 is required and sufficient for pathological growth. Cand2 knock-out mice are protected against pathological remodeling. Mechanistically, Cand2 post-transcriptionally controls the expression of G-protein coupled receptor 5 (Grk5) expression *in vitro* and *in vivo*, which in turn is linked to transcription of hypertrophic genes driven by myocyte enhancer factor 2 (MEF2).

Our data highlighted a novel mechanism where translational, mTORC1-dependent, control of Cand2 expression results in increased expression of Grk5 which results in transcriptional reprogramming in diseased cardiac myocytes (mRNA translation controls transcriptional activity). This linked mTOR dependent cytosolic signaling events, that drive specific mRNA translation, to transcriptional activity.

## Results

### Cand2 is a myocyte-specific protein translationally upregulated during pathological stress

To define mRNAs translationally regulated by mTOR signaling in response to pathological cell growth, cultures of neonatal rat ventricular cardiomyocytes (NRCMs) were acutely treated with the α-1 adrenoreceptor agonist phenylephrine (PE) and with Torin 1, which inhibits mTOR by binding to the ATP-binding site in the kinase domain (Thoreen *et al*, 2012). Specific inhibition of mTOR-dependent protein signaling by Torin 1 was confirmed by immunoblots (**EV1**). Ribosome profiling (Ribo-seq) has emerged as a quantitative technique to study global gene expression, and overcomes limitations of classical expression analysis, as it directly quantifies the number of translating ribosomes (Ingolia *et al*, 2009). The effect of Torin 1 on mRNA translation was analyzed by Ribo-seq in cultured cardiomyocytes (**Fig. 1A**). Total cellular RNA was also collected for parallel RNA-seq to quantify mRNA abundance (the full list is provided in **Source Data set**).

**Figure 1.**
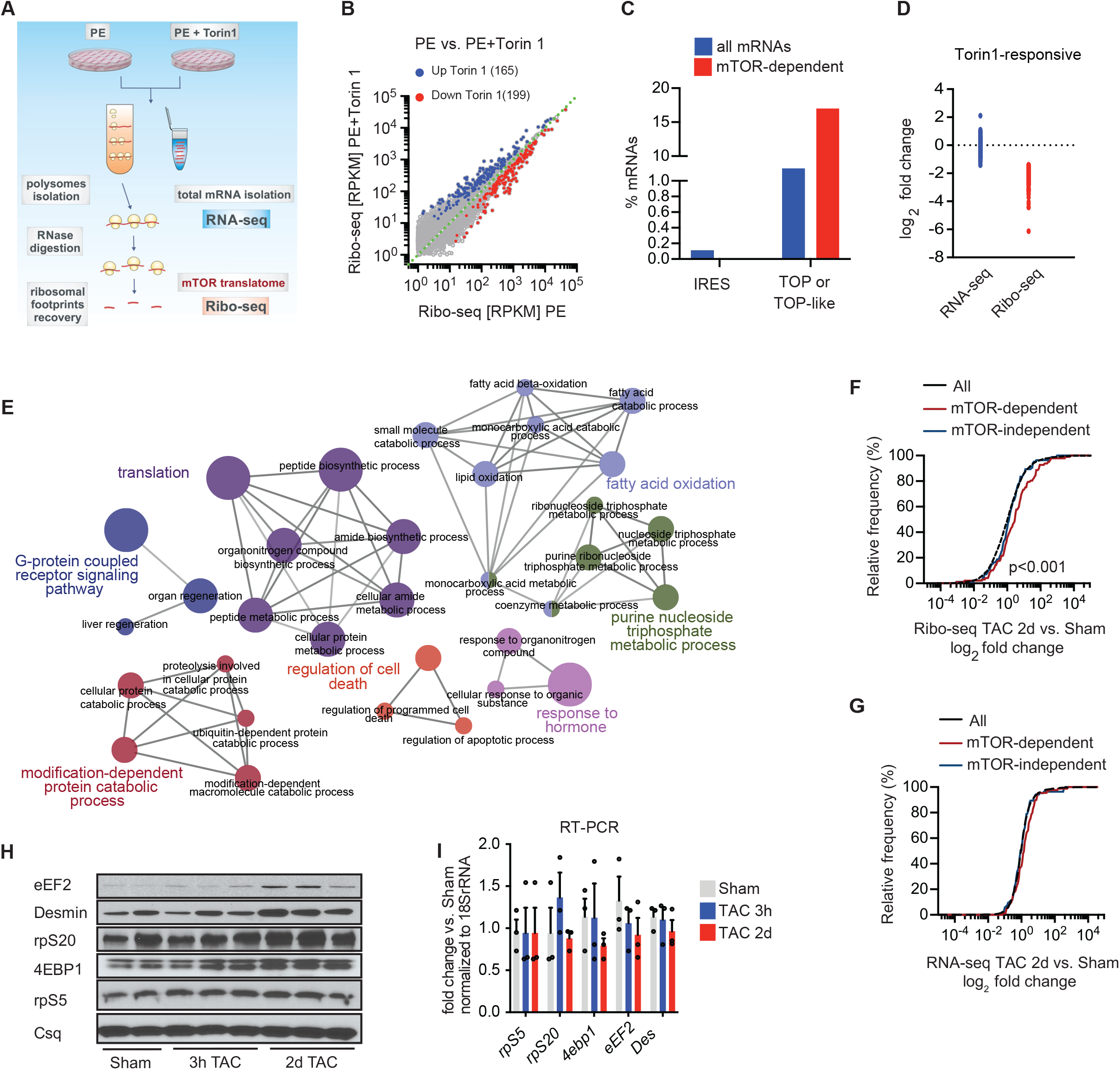
Identification of the mTORC1-dependent cardiac translatome. **(A)** Experimental design of translational profiling of mTOR-dependent mRNAs *in vitro* in NRCMs. Polysomal fractions were isolated in a sucrose gradient and ribosomal footprints were recovered by ribonuclease digestion. NRCMs were treated with vehicle or Torin1 to block mTOR pathway. **(B)** Scatter plot of ribosome occupancy in Ribo-seq of cultured cardiac myocytes in response to PE and Torin 1 treatment (blue dots - mRNAs downregulated after Torin 1, red dots - upregulated mRNA. **(C)** Percentage of TOP or TOP-like containing mRNAs in total and Torin1-sensitive mRNA pools. **(D)** Transcriptional (RNA-seq) vs. Translational (Ribo-seq) control of mTOR-regulated mRNAs. **(E)** Network of enriched GO categories (biological process) among mTOR-sensitive genes into several functional groups represented by circles of different colors. Each node represents one enriched GO category with the size of the node is proportional to the number of transcripts. Edges indicate similarity (between the two connected GO categories). **(F)** Cumulative fraction of mRNAs relative to their fold change of Ribo-seq and **(G)** RNA-seq (TAC vs. sham) between all transcripts mTOR-dependent and independent transcripts two days after TAC surgery. **(H)** Immunoblot and **(I)** RT-qPCR representing levels of proteins and corresponding mRNAs with defined 5’TOP motifs 3 hours and 2 days after TAC.

We identified 199 genes that decreased and 165 that increased (log2 FC > 1 or < −1) in Ribo-seq in an mTOR-dependent manner in response to PE treatment (**Fig. 1B**). Among the suppressed genes, 17% have a known TOP or TOP-like motif in the 5’ UTR (**Fig. 1C**), which is known to be sensitive to mTOR inhibition (Thoreen *et al*, 2012), and the suppressed genes are predominantly regulated on the translational level (**Fig. 1D**). Immunoblotting confirmed a decrease in the expression of selected mTOR-dependent genes such as eEF2 and rpS5 at the protein level upon Torin 1 treatment (**EV1**) but not at the transcript level. In contrast, molecular markers of pathological growth such as *Nppb* and *Nppa* were still induced (**EV1**). Stimulation of NRCMs with PE resulted in a strong increase in mRNAs in polysomal fractions (**EV1**). The increase of mRNAs in the polysomal fraction induced by PE was blocked by Torin 1, confirming that Torin 1 blocks the recruitment of ribosomes into polysomal fractions. The effects of inhibiting of mTOR on cardiac myocyte growth were examined in isolated cardiac myocytes stimulated with PE for 24h. Torin 1 inhibited PE-induced hypertrophy as assessed by cell surface area measurements (**EV2**). To measure the effect of Torin 1 on cap-dependent translation in cells, we used a dual-luciferase reporter vector that distinguishes cap-dependent versus cap-independent translation by separating Renilla luciferase from firefly with the Polio IRES(Poulin *et al*, 1998) (**EV2**). Therefore, the Renilla/firefly ratio would determine the cap-dependent translation ratio. Torin 1 inhibited cap-dependent, but not IRES-dependent translation measured by luciferase activity.

In line with the *in vitro* data, mTOR inhibition with Torin 1, *in vivo*, blocked pathologic growth induced by acute transverse constriction (TAC) surgery, a common model for cardiac hypertrophy and remodeling (**EV2**) (Doroudgar *et al*, 2019). Thus, mTOR-dependent and cap-dependent protein synthesis are increased during hypertrophic growth and this is necessary for induction of cell growth, *in vitro* and *in vivo.*

Gene ontology (biological processes) analysis of the genes that were suppressed by Torin 1 showed strong enrichment for those involved in translation, metabolism, as well as signaling cascades (**Fig. 1E**). Overall, this dataset defined a specific subset of genes that are regulated in an mTOR-dependent manner in response to neurohumoral stimulation with PE. Next, we followed our identified mTOR-dependent genes in a previously published *in vivo* data set after TAC surgery (Doroudgar *et al*, 2019). mTOR-dependent genes were translationally upregulated in response to TAC surgery (**Fig. 1F-G**). Increased expression of identified mTOR targets independent from changes of transcript levels such as eEF2, Desmin, rpS5, or rpS20 were confirmed using immunoblots and RT-qPCRs from heart lysates after TAC surgery (**Fig. 1H-I**).

Among the 199 genes with decreased translation upon Torin 1 treatment, 17 were significantly increased in the *in vivo* data set after TAC surgery (Doroudgar *et al*, 2019) (**Fig. 2A**). Among this subset of genes that were increased by pressure overload *in vivo* and dependent on mTOR activity *in vitro*, was Cand2, a muscle specific protein with unknown role in the heart ^12^. (**Table Fig. 2A**). Since its role in cardiomyocytes was completely unknown, we aimed to characterize the role of Cand2 during pathological remodeling. Torin 1-induced downregulation of Cand2 translation was not due to transcriptional regulation assessed by parallel RNA-seq (**Fig. 2B**). mTOR activity increased Cand2 protein levels *in vitro*, as well, but had no impact on its transcript levels measured by RT-qPCR (**Fig. 2C-D**), confirming the results from the Ribo-seq data sets. Ribo-seq data revealed an increased expression of Cand2 two days after TAC (**Fig. 2E**). In contrast, Cand2 translation was unchanged in early (3 hours after TAC) and chronic pressure overload (2 weeks after TAC) (**Fig. 2F-G**). Translational upregulation of Cand2 in the heart was also detected on the protein level when compared by immunoblotting to sham-operated mice (**Fig. 2H**). Increased Cand2 levels *in vivo* were also not caused by increased Cand2 transcript levels as shown by RT-qPCR (**Fig. 2I**). To assess whether Cand2 expression is controlled by mTORC1 *in vivo*, Torin 1 was injected in mice after TAC or sham surgery and Cand2 levels were assessed by immunoblotting (**Fig. 2H**). The known mTORC1-sensitive 40S ribosomal protein 5 (rpS5) was used as a positive control (Uprety *et al*, 2018). Along with rpS5 inhibition by Torin 1, increased Cand2 protein levels after TAC surgery were completely blocked by Torin 1, suggesting that Cand2 mRNA belongs to mTORC1-sensitive genes, *in vivo*, as well. Since mTORC1 specifically regulates translation of mRNAs with 5’TOP motifs, we placed the 5’ UTR of Cand2 mRNA upstream of *Renilla* luciferase to investigate its regulatory potential in a mTORC1-dependent manner (**Fig. 2J**). Cand2 5’ UTR showed a similar degree of reporter inhibition with Torin 1 treatment as the known TOP motif-containing 5’ UTR of eEF2. This confirmed mTORC1-dependent translation of Cand2 and suggests the presence of a regulatory motif in its 5’ UTR. Taken together, mTORC1 controls the expression levels of Cand2 in response to pathological stimulation both *in vitro* and *in vivo*.

**Figure 2.**
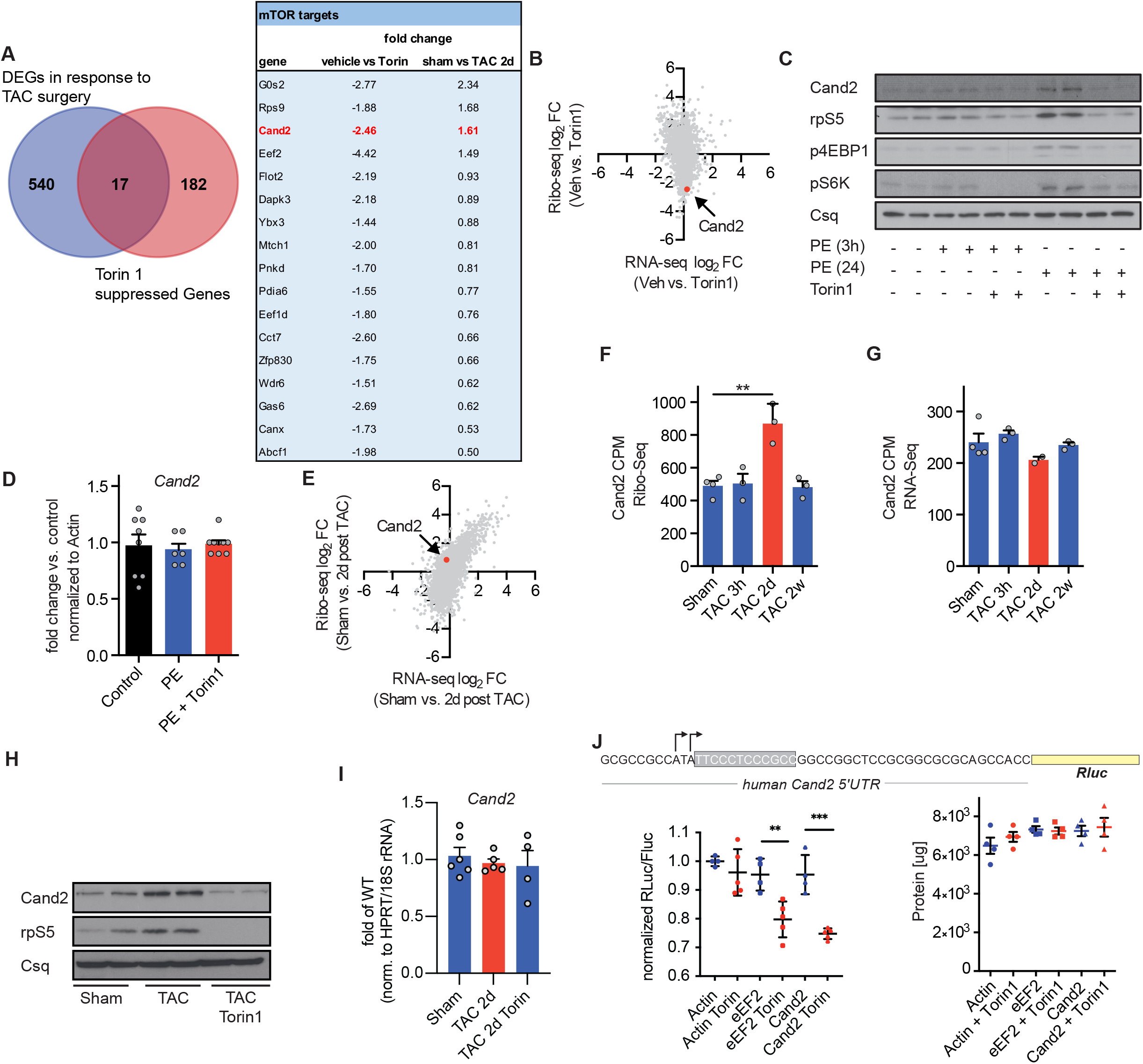
Cand2 is translationally upregulated during cardiac pathological stress. **(A)** Venn diagram showing a group of transcripts translationally upregulated *in vivo* 2 days after TAC and regulated by mTORC1 *in vitro* in NRCMs treated with Torin1. Table with top the most translationally regulated mTOR-dependent transcripts *in vitro* after Torin1 treatment of NRCMs and *in vivo* after TAC **(B)** Scatter plot of Ribo-seq vs. RNA-seq upon NRCMs treatment of Torin1 shows Cand2 among translationally downregulated mRNAs. **(C)** Protein expression of Cand2 in NRCMs treated with PE for 3 and 24 h, and 24h of Torin1 treatment. mTORC1 induction was monitored by downstream targets, p4EBP1 and pS6K proteins. rpS5 - example of mTOR-dependent mRNA used as a positive control. **(D)** Cand2 mRNA levels in NRCMs treated with PE and Torin1 measured by RT-qPCR. **(E)** Scatter plot of Ribo-seq vs. RNA-seq 2 days after TAC showing an increase of Cand2 translation **(F)** Ribo-seq and **(G)** RNA-seq expression data of Cand2 in sham and TAC operated animals (CPM - count per million). **(H)** Immunoblots of Cand2 expression in left ventricles 2 days post-TAC and after mice injection with Torin1. rpS5 used as a positive control of mTOR-dependent mRNA. **(I)** Cand2 mRNA 2 days after TAC measured by RT-qPCR. **(J)** Schematic representation of reporter consisting of human Cand2 5’UTR and downstream *Renilla* luciferase coding sequence. Potential TOP-like motif is highlighted in a grey box. Grey arrows indicate transcription start sites in the adult heart according to Database of Transcriptional Start Sites (Release 10.1). Cand2 5’UTR effect on translation measured by luciferase activity in normal conditions and after mTORC1 block with Torin1. eEF2 - positive control of reporter with defined 5’TOP motif and non-TOP 5’UTR of β-Actin used as a negative control. Total protein content in lysates used for luciferase assay.

To further characterize Cand2, we analyzed its expression profile in different organs in mice. Consistently with a previous report (Aoki *et al*, 1999), Cand2 protein was expressed predominantly in muscle tissues and was highly abundant in the left ventricle (**Fig. 3A**). Quantitative RT-PCR analysis also revealed the highest Cand2 mRNA level in the heart (**Fig. 3B**). Immunofluorescent staining confirmed specific Cand2 expression only in cardiomyocytes (**Fig. 3C**), suggesting that Cand2 protein expression is restricted to muscle cells in the heart. Subsequently, we analyzed Cand2 subcellular localization in isolated cardiomyocytes by immunofluorescent staining (**Fig. 3D**). Consistently with studies on skeletal muscle cell maturation (Shiraishi *et al*, 2007), endogenous Cand2 was located in the nucleus and cytoplasm in cardiac myocytes. Subcellular fractionation of the left ventricle confirmed the cytoplasmic and nuclear distribution of Cand2 (**Fig. 3E**).

**Figure 3.**
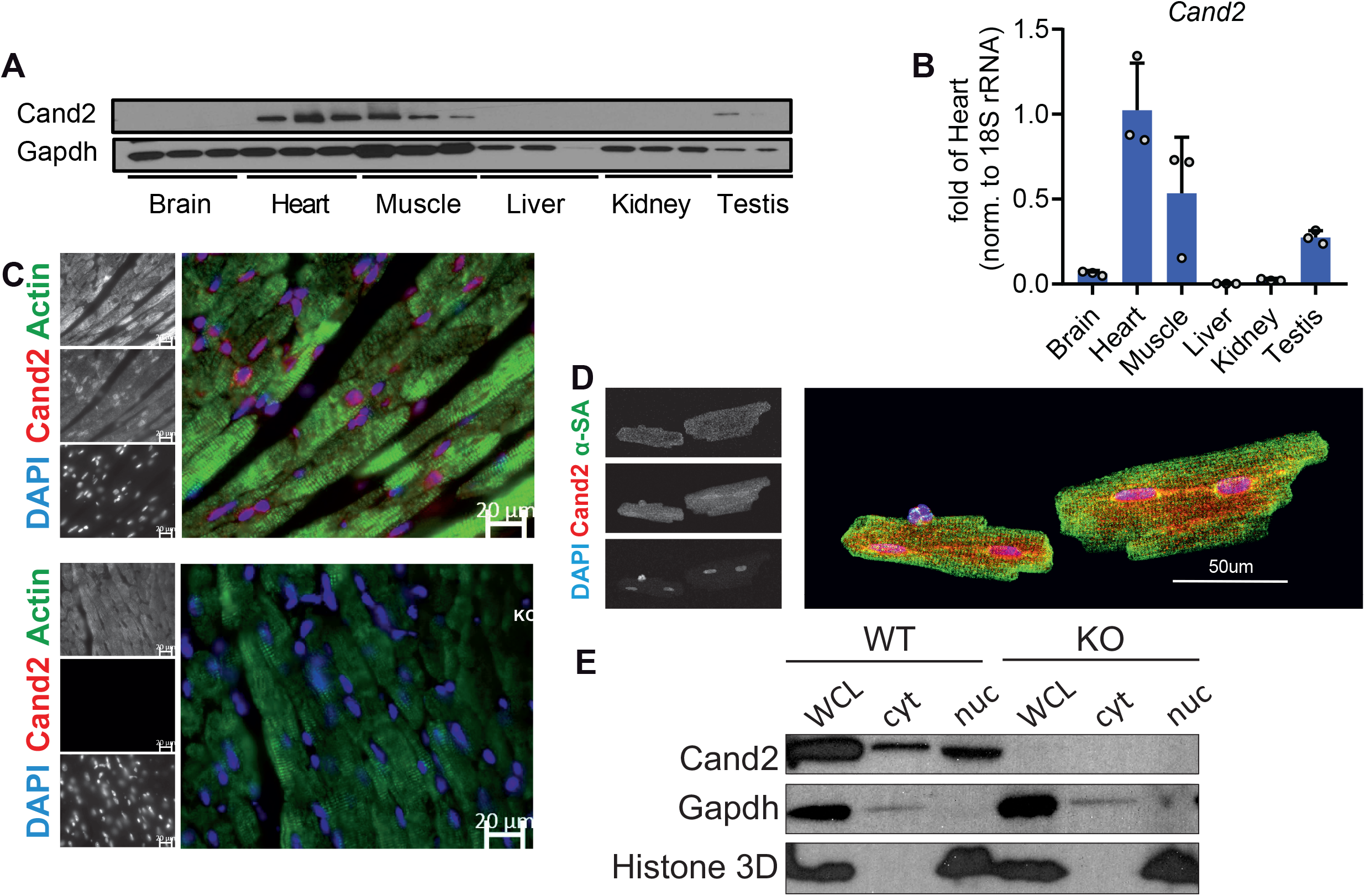
Cand2 is a muscle-specific protein. **(A)** Representative immunoblot of Cand2 protein levels in mouse organs. **(B)** Cand2 expression in mouse organs measured by RT-qPCR. **(C)** Immunofluorescent staining of Cand2 (red), actin (green) and nuclei (DAPI, blue) in paraffin sections from left ventricles of WT and Cand2 KO mice. Scale bar 20μm **(D)** Immunofluorescence of adult rat cardiomyocytes of Cand2 (red), sarcomeric actin (green), and nuclei (blue). Scale bar 50μm. **(E)** Immunoblot of subcellular fractionation of left ventricle (LV). Gapdh and Histone 3D (H3D) used as cytoplasmic and nuclear markers, respectively. WCL - whole cell lysate, cyt - cytoplasmic fraction, nuc - nuclear fraction.

### Cand2 contributes to cardiomyocytes pathological growth *in vitro* and *in vivo*

Next, we studied the effect of Cand2 on cardiomyocyte cell size after Cand2 knock-down or overexpression (**Fig. 4A-B**). Cand2 depletion *in vitro* decreased cell surface area and blunted the response to neurohormonal stimulation with PE treatment (**Fig. 4C-D**). In contrast, Cand2 overexpression induced cardiomyocyte growth, and the increase in cell size after overexpression was comparable to control cells after PE (**Fig. 4E**). *Nppa* levels correlated to Cand2 expression: they were decreased when Cand2 was depleted and elevated in Cand2 overexpression in untreated and PE stimulated cardiomyocytes (**Fig. 4F**).

**Figure 4.**
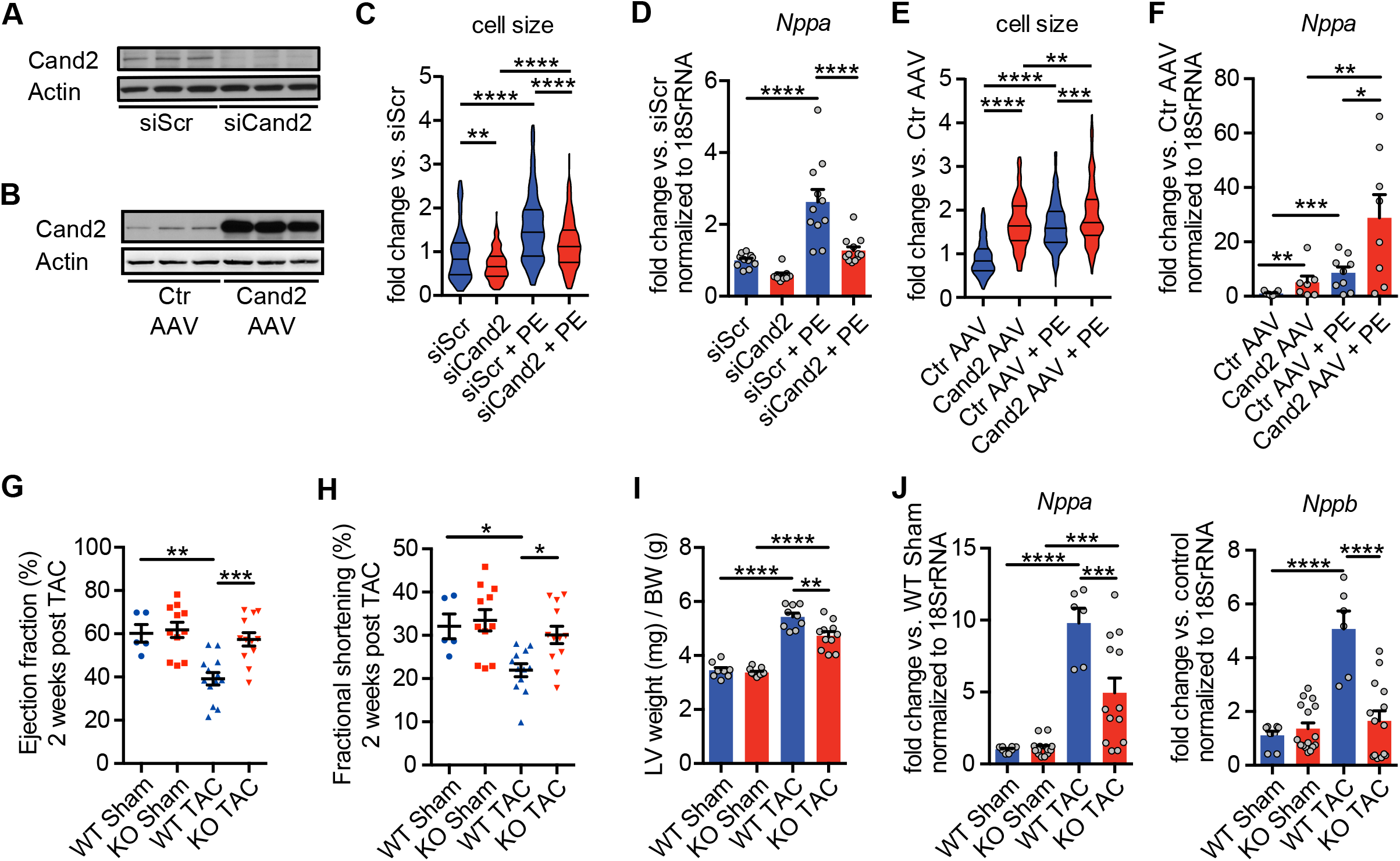
Cand2 is required for cardiac hypertrophy induction. **(A)** Representative immunoblot of Cand2 KD in NRCMs with Cand2 specific siRNA (siScr - control scrambled siRNA). **(B)** Cand2 OE in NRCMs (Cand2 AAV6; Ctr AAV – control AAV6 vector) analyzed by Western blot. β-Actin – housekeeping protein. **(C)**. Cell surface area measurement of NRCM after Cand2 KD and 24h PE stimulation **(D)**. RT-qPCR analysis of *Nppa* transcript level after Cand2 KD. **(E)** NRCMs size measurement after Cand2 OE and 24h PE stimulation. **(F)** *Nppa* mRNA levels *in vitro* after Cand2 OE and PE treatment measured by RT-qPCR. **(G)** Ejection fraction 2 weeks after TAC in WT and Cand2 KO mice measured by echocardiography. **(H)** Fractional shortening 2 weeks after TAC in WT and Cand2 KO mice measured by echocardiography. **(I)** LV weight to body weight ratio of WT and Cand2 KO mice subjected to sham and TAC surgeries. **(J)** *Nppa* and *Nppb* mRNA levels 4 weeks after TAC measured by RT-qPCR. Error bars indicate means ± SEM; * - *P* ≤ 0.05, ** - *P* ≤ 0.01, **** - *P* ≤ 0.0001

To study the role of Cand2 *in vivo*, a novel Cand2 knock-out (KO) mouse was generated (**EV3**). Cand2 deficient mice were viable and cardiac function as assessed by ejection fraction, as well as left ventricle weight to body weight ratio remained unchanged compared to littermate control animals (**EV3**). In line with this, Cand2 KO did not alter the levels of fetal gene markers normally upregulated in the adult heart during stress, such as *Nppa* and *Nppb* (**EV3**). Moreover, Cand2 knock-out did not disturb the levels of Cand1 (**EV3**)

Next, Cand2 KO mice along with wild-type (WT) littermates were subjected to TAC or sham surgery and phenotypic consequences were analyzed 4 weeks after surgery. TAC operation resulted in decreased ejection fraction, fractional shortening and increased LV/BW ratio in WT mice (**Fig. 4G-I**). In contrast, Cand2 deficient mice showed preserved cardiac function as well as significantly decreased LV/BW ratio compared to TAC-operated WT mice. Moreover, TAC-induced *Nppa* and *Nppb* levels in WT mice were 1.9-fold and 3-fold reduced in Cand2 KO mice, respectively (**Fig. 4J**). Overall, these results suggest that Cand2 deletion protects against pathological remodeling in response to pressure overload.

### Cand2 controls Grk5 expression and affects MEF2-dependent transcription

Next, we performed transcriptome profiling of left ventricles from wild-type and Cand2 knock-out mice to identify targets that might be regulated by Cand2. mRNA sequencing identified only small changes in the transcriptome when Cand2 was deleted (**Fig. 5A**). Interestingly, pathway analysis revealed that most of the Cand2-dependent genes are enriched in categories regulating transcriptional control and proteasomal function. Among the Cand2-dependent group of genes involved in transcriptional control, the G-protein coupled receptor kinase 5 (Grk5) has been one of the most regulated genes in Cand2 knock-out mice (**Fig. 5A**). While the canonical role of Grks is to phosphorylate agonist bound G-protein coupled receptors, which promotes the binding of an arrestin protein to the receptor, resulting in subsequent desensitization of the receptor, Grk5 has been found to localize to the nucleus *via* a nuclear localization sequence. Previous work showed that following pressure overload, Grk5 accumulates in the nucleus of cardiomyocytes and acts as a class II histone deacetylase (HDAC) kinase, phosphorylating HDAC5 specifically, leading to its nuclear export and de-repression of the transcription factor MEF2 (Zhang *et al*, 2011).

**Figure 5.**
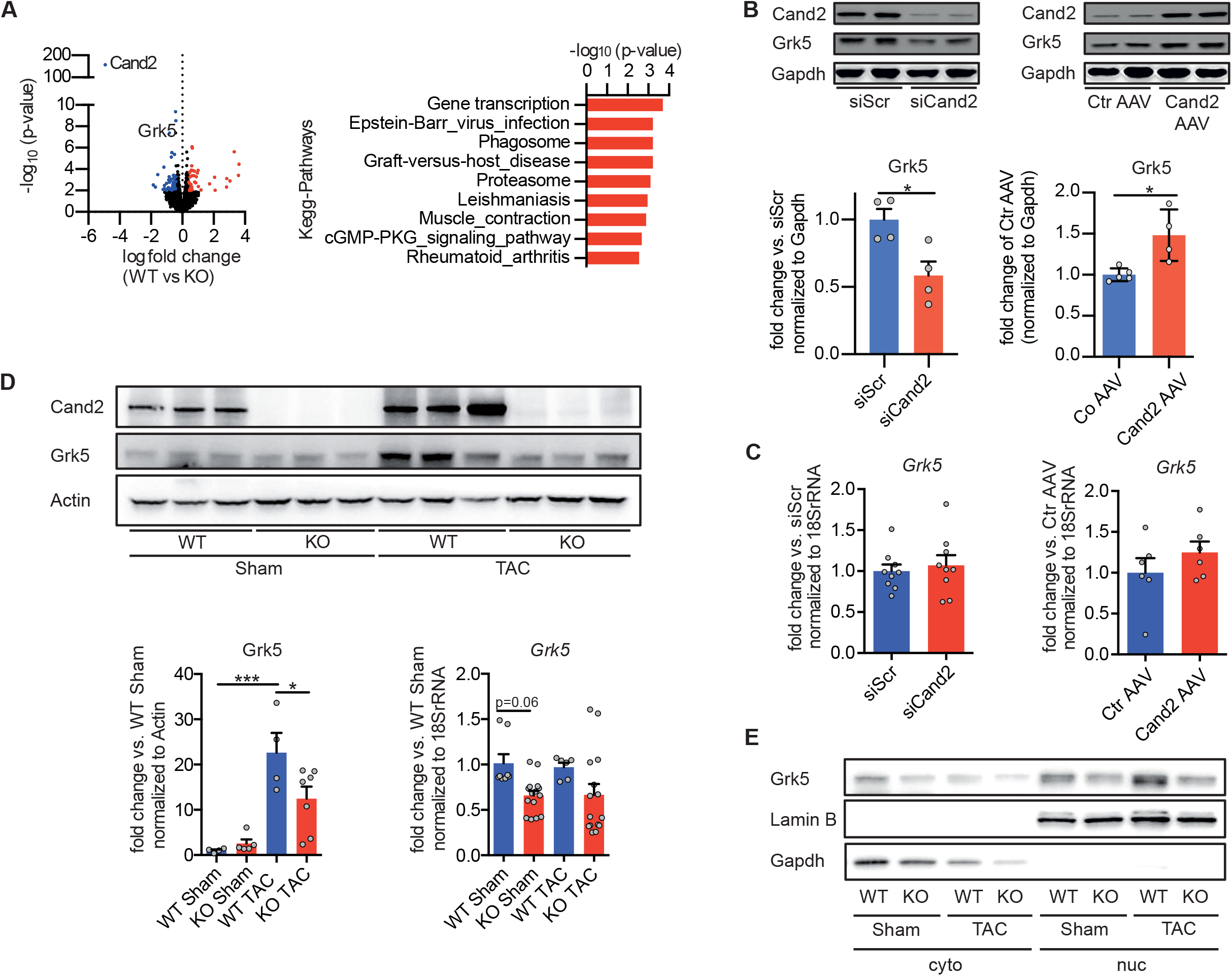
Cand2-dependent Grk5 expression. **(A)** Enrichment of Kyoto Encyclopedia of Genes and Genomes (KEGG) terms for differentially expressed genes in LV of Cand2 KO vs. WT mice (right panel). Volcano plot of up – (RNA-seq log2-fold change of count per million >1; red dots) and downregulated (>1; blue dots) transcripts in LV after Cand2 KO (left plot). **(B)** Immunoblots and gel quantification of Grk5 protein levels *in vitro* after Cand2 KD with siRNA and Cand2 OE with AAV6. **(C)** Grk5 expression levels in NRCMs depleted of and overexpressing Cand2 measured by RT-qPCR. **(D)** Grk5 protein levels *in vivo* in LVs of WT and Cand2 KO mice 4 weeks after sham and TAC analyzed by immunoblotting (upper panel) and its quantification (left bottom graph), and Grk5 mRNA levels in indicated conditions (right bottom graph). **(E)** Representative immunoblot of Grk5 protein in cytoplasmic and nuclear fractions from left ventricle lysates of WT and Cand2 KO mice after TAC surgery. Lamin B – nuclear marker, Gapdh – cytoplasmic marker.

First, we analyzed Grk5 expression after gain- or loss-of-function of Cand2 *in vitro* in neonatal rat ventricular myocytes (NRCMs) (**Fig. 5B-C**). Cand2 overexpression increased Grk5 protein in NRCMs but did not change its mRNA levels. In contrast, Grk5 protein was downregulated in Cand2 knock-down and Grk5 transcript level again remained unaffected. Next, we further analyzed levels of Grk5 in Cand2 KO mice in TAC-induced pressure overload 2 weeks after surgery (**Fig. 5D**). Quantitative RT-PCR analysis did not reveal any significant changes in Grk5 mRNA levels after TAC in Cand2 KO mice (**Fig. 5D**). In line with other reports, Grk5 protein was highly increased in TAC-operated wild-type mice. Lack of Cand2 led to an approximately 10-fold decrease of Grk5 protein levels after TAC. Since Grk5 has been shown to translocate to the nucleus of cardiomyocytes following pressure overload *via* its nuclear localization signal (NLS) (Johnson *et al*, 2004),(Martini *et al*, 2008),(Gold *et al*, 2013), we examined Grk5 levels in the cytoplasm and nucleus *in vivo* in Cand2 KO mice (**Fig. 5E)**. Neither TAC nor lack of Cand2 affected Grk5 levels in the cytoplasmic fraction, but Grk5 levels increased in the nucleus in WT mice in response to TAC, which was blocked in Cand2 mice.

Since Grk5 regulates the activity of the transcription factor MEF2, we analyzed MEF2 activity by using a luciferase reporter assay. Cand2 knock-down decreased MEF2 luciferase activity by 3.5-fold *in vitro* (**Fig. 6A**). Moreover, the expression of direct MEF2-dependent mRNAs such as *Nr4a1* and *Xirp2* (Lehmann *et al*, 2018),(Huang *et al*, 2006) were downregulated when Cand2 was depleted (**Fig. 6B**). Conversely, the levels of *Nr4a1* and *Xirp2* increased significantly in NRCMs overexpressing Cand2 (**Fig. 6C**). Inline, levels of *Xirp2* increased in WT mice in response to TAC but were blocked in Cand2 KO mice (**Fig. 6D**). Similarly, mTOR inhibition with Torin1 resulted in decreased Grk5 protein levels and MEF2 luciferase activity in NRCMs, without changes in mRNA levels of *Grk5* (**Fig. 6E**). Next, we confirmed decreased Cand2 and Grk5 protein but not transcript levels after depletion of Raptor, the essential component of mTORC1, in myocytes (**Fig. 6F**). Raptor knock-down caused decreased protein levels of Cand2 and Grk5 (**Fig. 6G**). Moreover, RT-qPCR analysis revealed downregulation of direct MEF2 targets such as *Nr4a1* and *Xirp2* as well as *Nppa* after knock-down of Raptor (**Fig. 6H**), suggesting that Grk5 expression and Grk5-mediated MEF2 activity are under direct mTORC1 control. Overall, these results suggest that Cand2 dependent regulation of Grk5 levels is associated with increased MEF2 transcriptional activity and similarly to Cand2, Grk5-MEF2 is downstream of mTORC1 as well.

**Figure 6.**
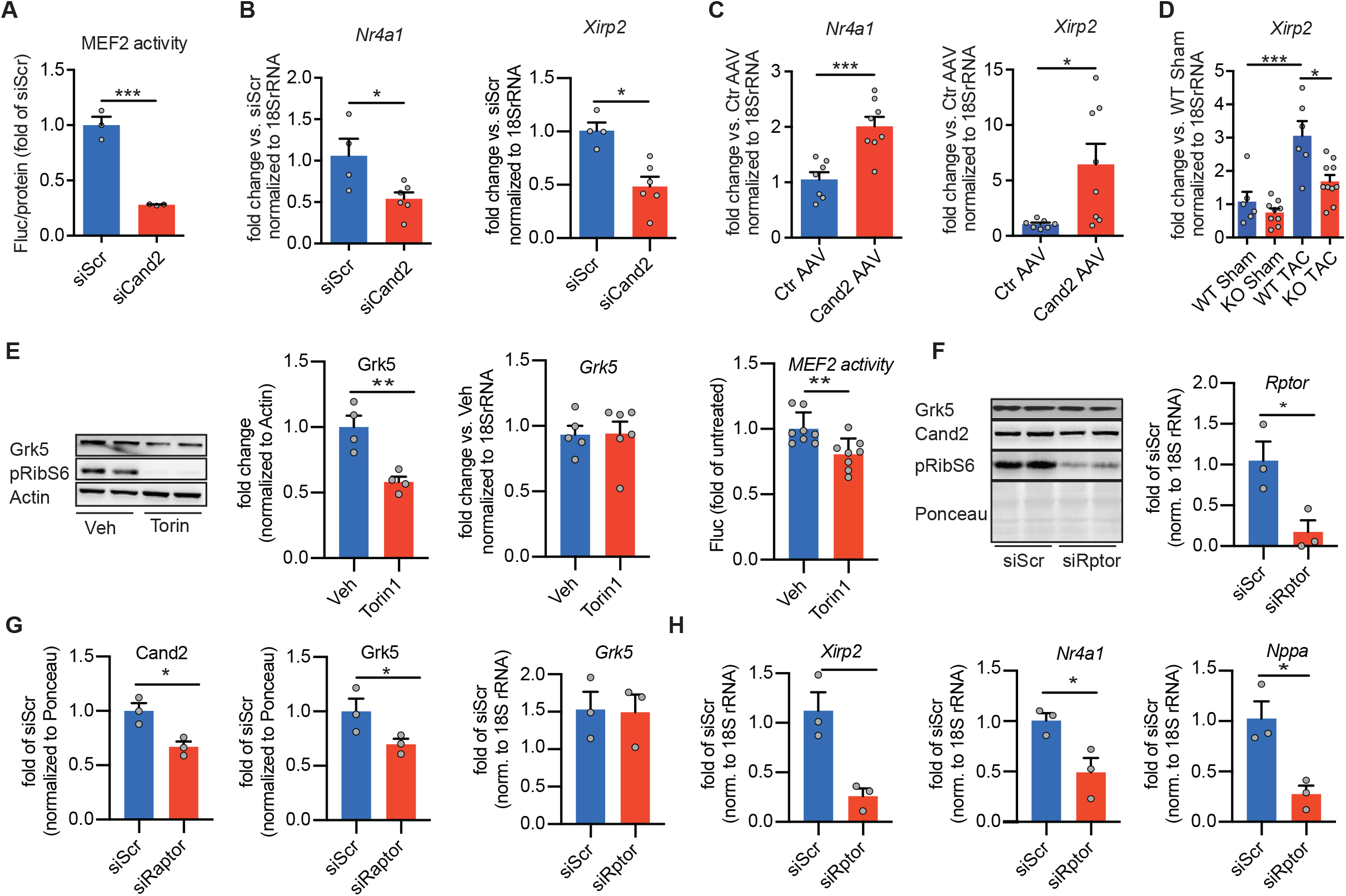
Cand2 regulates MEF2 transcriptional activity. **(A)** Luciferase based quantification of MEF2 activity after Cand2 KD in NRCMs. **(B)** mRNA levels of MEF2-dependent genes *Nr4a1* and *Xirp2* in NRCMs with Cand2 KD. **(C)** mRNA levels of MEF2-dependent genes *Nr4a1* and *Xirp2* in NRCMs with Cand2 OE. **(D)** RT-qPCR analysis of MEF2 targets in LVs of WT and Cand2 KO mice after sham and TAC surgeries. **(E)** Grk5 protein, mRNA levels in NRCMs and MEF2-luciferase activity after 24h of Torin 1 treatment. **(F)** Cand2 and Grk5 protein levels and Rptor transcript level in NRCMs after Raptor KD analyzed by Western blot **(G)** Quantifications of Cand2 and Grk5 protein levels and Grk5 transcript level in NRCMs depleted of Raptor. **(H)**, *Nppa*, and MEF2 targets *Nr4a1* and *Xirp2* mRNA levels in NRCMs after Raptor KD. Error bars indicate means ± SEM; * - *P* ≤ 0.05, ** - *P* ≤ 0.01, *** - *P* ≤ 0.001, **** - *P* ≤ 0.0001

### Cand2 regulates Grk5 protein level by Cullin1 neddylation inhibition

Our findings showed that Cand2 depletion results in decreased Grk5 protein levels independent from changes in transcript levels (**Fig. 5B**), suggesting post-transcriptional regulation of Grk5 expression by Cand2. Cand proteins modulate the activity of Cullin–RING E3 ubiquitin ligases (CRLs). The activity of CRLs is largely dependent on Cullin neddylation, whereby a covalent modification of Cullin1 with the ubiquitin-like protein Nedd8 will induce the conformational rearrangement of Cullin1. Cand2 protein has been shown to interact with and sequester unneddylated Cullin1. To find out whether Cand2, Cullin1, and Grk5 directly interact in cardiomyocytes, co-immunoprecipitations were performed. Endogenous Cullin1 as well as Grk5 was detected after Cand2 immunoprecipitation, whereas Cul4 did not bind Cand2 (**EV4**). Additionally, we performed proximity ligation assay (PLA) in neonatal cardiomyocytes isolated from wild type and Cand2 knock-out mice (**EV4**). PLA signal was detected in cardiomyocytes expressing endogenous Cand2, but not in KO cells, suggesting that Cand2 is associated with Grk5 specifically. Although both proteins were observed ubiquitously in the cell, PLA signal was concentrated in the nucleus. These results support the idea that Cand2 interacts in a complex with Cullin1 with Grk5.

Cand proteins selectively bind to the unnedyladed pool of Cullin1, thus affecting SCF ubiquitin ligase complex (Liu *et al*, 2002). Thus, we sought to determine the effect of Cand2 on the overall neddylation status of Cullin1 in cardiomyocytes overexpressing and depleted of Cand2 (**Fig. 7A and B**). Neddylated Cullin1 migrates slower during electrophoresis and can, therefore, be distinguished from unnedylated Cul1. Manipulation of Cand2 expression affected only the neddylated Cul1 levels, whereas the unnedylated fraction remained unchanged. The amount of neddylated Cul1 was slightly but significantly increased when Cand2 was silenced and reduced in Cand2 overexpressing cells, suggesting that Cand2 may regulate Cul1 activity (**Fig. 7A-B**).

**Figure 7.**
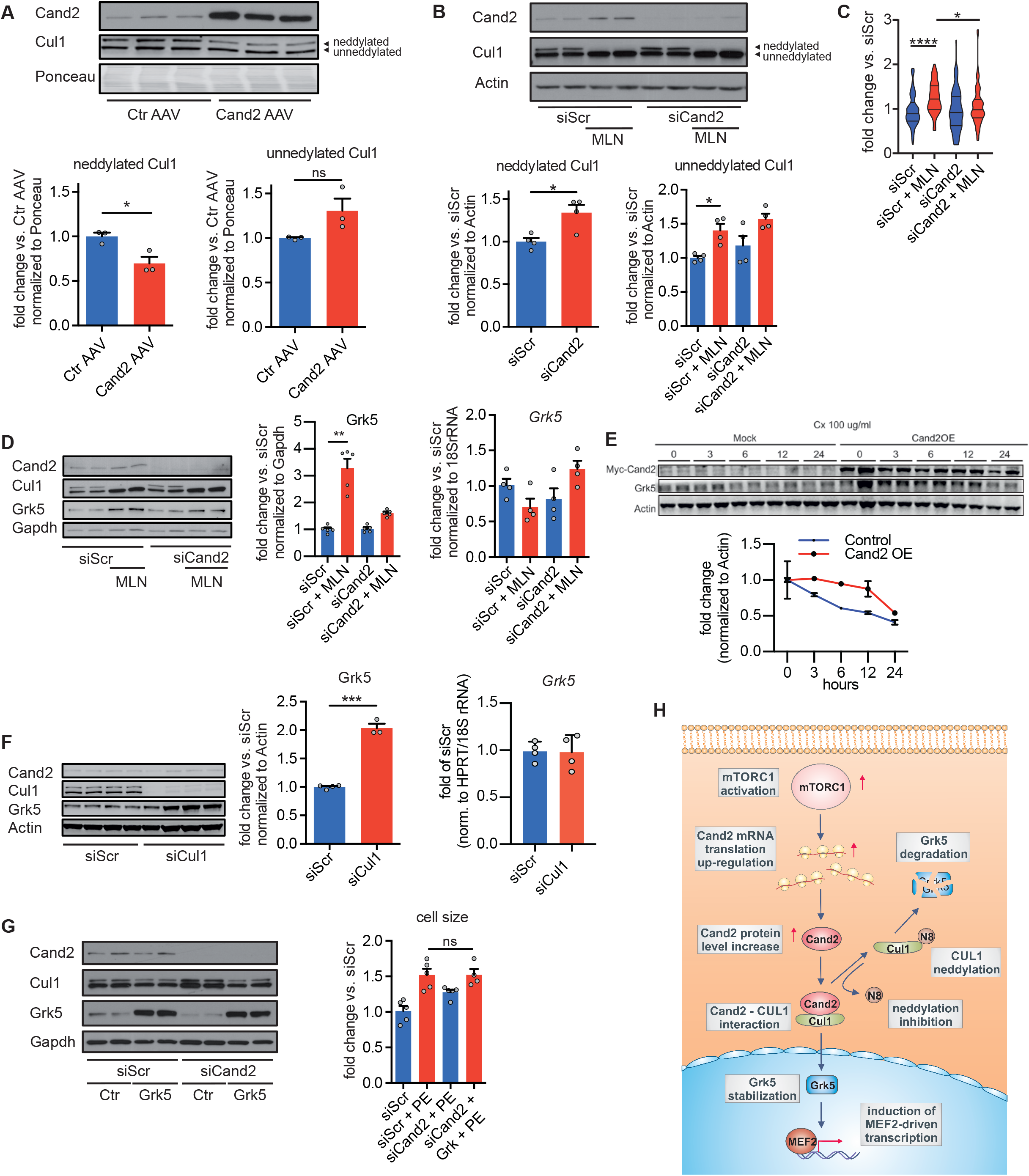
Cand2 and Cullin1 neddylation modulate Grk5 protein level. **(A)** Representative immunoblot and its quantification of protein levels of neddylated and unnedylated Cul1 in NRCMs after Cand2 OE. **(B)** Cul1 neddylation analyzed by Western blot in NRCMs depleted of Cand2. NRCMs were treated with MLN4924 to block Cul1 neddylation. The graphs show immunoblot quantification. **(C)** Cell size area measurement of NRCMs after Cand2 KD and neddylation inhibition (MLN). **(D)** Grk5 expression in NRCMs upon Cand2 KD and Cul1 neddylation inhibition examined by immunoblot and RT-qPCR. **(E**) The half-life of Grk5 in HeLa cells overexpressing Cand2 measured by cycloheximide chase. **(F)** Grk5 protein and mRNA levels *in vitro* after Cul1 KD and immunoblot quantification. **(G)** Comparison of NRCMs size with Cand2 KD, Grk OE after PE stimulation. Immunoblot confirms Cand2 KD and Grk5 OE. Error bars indicate means ± SEM; * - *P* ≤ 0.05, ** - *P* ≤ 0.01, *** - *P* ≤ 0.001, **** - *P* ≤ 0.0001 **(H)** Model of Cand2-dependent stabilization of Grk5 by Cul1 neddylation inhibition in mTOR-dependent manner.

In line with previous reports (Zou *et al*, 2019), inhibition of neddylation by using a specific inhibitor (MLN4924) was sufficient to induce cardiomyocyte hypertrophy, which was completely blocked by Cand2 knock-down **(Fig. 7C)**. Interestingly, we found that MLN4924 mediated inhibition of Cul1 neddylation alone was sufficient to induce an approximately 2.5- fold increase of Grk5 protein levels, independent from transcript levels measured by RT-qPCR **(Fig. 7D)**. The deneddylation of Cul1 by MLN4924 led to approximately 2.5-fold increase of Grk5 protein, independently from transcript levels measured by RT-qPCR. This induction was largely attenuated after Cand2 knock-down, indicating that Cand2 is required for the induction of Grk5 after neddylation inhibition **(Fig. 7D).** Additionally, neddylation inhibition did not affect Cand2 transcript levels (**EV5**). Neither Cand2 knock-down nor overexpression influenced Cul1 mRNA levels, indicating that Cand2 interacts with Cul1 on the protein level (**EV5**). To show that Cand2 regulates Grk5 protein stability, we assessed Grk5 half-life in cycloheximide chase assay (**Fig. 7E**). The Grk5 protein half-life was prolonged from approximately 18 to 27 hours when Cand2 was overexpressed suggesting that Grk5 is stabilized by Cand2.

Similar to neddylation inhibition, knock-down of Cul1 increased Grk5 protein amount by 2-folds (**Fig. 7F**). This strongly suggests that Grk5 degradation is mediated by Cullin-RING E3 ubiquitin ligase complex. Quantitative RT-PCR analysis revealed that all changes in Grk5 levels caused by MLN4924 treatment or Cand2/Cul1 silencing resulted independent from transcriptional changes (**Fig. 7D-F**).

Finally, to confirm that Cand2 affects cell size through Grk5, we examined whether Grk5 re-expression is sufficient to rescue the decreased cell size phenotype of Cand2-deficient cardiomyocytes. Grk5 overexpression was confirmed by immunoblot and was also functional in Cand2 knock-out cells **(Fig. 7G)**. While Cand2 knock-down again attenuated the hypertrophic growth response of cardiomyocytes, Grk5 overexpression was sufficient to restore hypertrophic cell growth to similar levels of control cardiomyocytes treated with PE. Taken together, we suggest a novel myocyte-specific and mTORC1-mediated signaling cascade. mTORC1-dependent, control of Cand2 expression results in activation of the pro-hypertrophic Grk5-HDAC4-MEF2 axis *via* post-transcriptional regulation of Grk5 protein levels by the Cul1-SCF ubiquitin ligase complex **(Fig. 7H)**.

## Discussion

This study provides evidence for a novel role of Cand2, a component of the SCF (SKP1-Cul1-F-box protein) E3 ubiquitin ligase complex, in the myocardium. Cand2 is a muscle-specific protein that we identified in a cardiomyocyte genome-wide screen for mTORC1-dependent gene expression control at the level of mRNA translation. Several studies and our own work showed that mTORC1-signaling controls physiological as well as pathological myocardial remodeling (Sciarretta *et al*, 2018; Völkers *et al*, 2014, 2013). The central role of mTORC1 in integrating multiple intra- and extracellular parameters requires an elaborate control system to accomplish fine-tuning of signaling to downstream targets. Clinically, targeting mTORC1 with rapamycin is already an established application to prevent re-stenosis after percutaneous coronary stent implantation. Systemic pharmacological or genetic mTORC1 inhibition prevented pathological hypertrophy and improved cardiac function in murine disease models (Buss *et al*, 2009; Shioi *et al*, 2003; Völkers *et al*, 2013). Still, the identity of mTORC1 translationally regulated mRNAs in the diseased heart were largely unknown.

Cand2 is a translationally upregulated gene dependent on the activity of mTORC1 during pathological stress both *in vitro* and *in vivo*, whereas Cand2 expression is not regulated during physiological stress. We found that Cand2 is required for pathological hypertrophy in cardiomyocytes by upregulation of the kinase Grk5 which is associated with the induction of pathological gene expression profiles. Indeed, Cand2 knock-out mice were protected against pathological remodeling, and cardiac dysfunction, which was associated with reduced levels of Grk5 and MEF2-dependent gene expression profiles. Conversely, overexpression of Cand2 was sufficient to drive pathological hypertrophy and gene expression *in vitro.* Cand2 expression was also found to be upregulated in human cardiomyopathy patients (van Heesch *et al*, 2019).

Cand2 was initially identified as a muscle-specific homolog of Cand1 and its only described function was the acceleration of myogenesis in skeletal myoblasts during differentiation (Aoki *et al*, 1999; Shiraishi *et al*, 2007). Interestingly, Cand2 has been linked to atrial fibrillation susceptibility using expression quantitative trait loci mapping, suggesting that it might play an important role in atrial myocytes as well. Increased levels of Cand2 correlated with a higher incidence of atrial fibrillation (Sinner *et al*, 2014; Wei *et al*, 2016; Gregers *et al*, 2017), suggesting that Cand2 might drive pathological remodeling in the atrium.

Mechanistically, Cand proteins modulate the activity of Cullin–RING E3 ubiquitin ligases (CRLs). The substrate-binding domain of Cullins, through their adaptors, can recruit hundreds of known substrate receptors that specifically target an even larger array of substrates. It has become increasingly evident that the cardiac ubiquitin-proteasomal function is not only highly dynamic but is also critical for healthy myocardium (Drews & Taegtmeyer, 2014). Just as CRLs regulate protein levels and function by covalent modification, they are subjected to modifications at the posttranslational level, which in turn regulate ligase activity. NEDD8, a ubiquitin-like protein, is covalently attached to a conserved lysine residue in the C-terminal Cullin homology domain. The activity of CRLs is largely dependent on Cullin neddylation, whereby Nedd8 induces conformational rearrangement of Cullins. Cand proteins have been shown to interact with and sequester unneddylated Cullins (Liu *et al*, 2018). Thus, Cand proteins have been initially characterized as inhibitors of Nedd8 conjugation, which initially implied that Cand proteins are negative regulators of CRLs. However, subsequent studies of Cand1 deficient cells and organisms suggested also a positive regulatory function of Cand on CRLs (Liu *et al*, 2018). Importantly, neddylation homeostasis is crucial for the integrity of the heart. Complete loss of neddylation in cardiac myocytes resulted in dilated cardiomyopathy with premature death (Su *et al*, 2013, 2011). Pharmacological inhibition of neddylation using a selective neddylation inhibitor MLN4924 promoted pathological hypertrophy and resulted in cardiac failure (Zou *et al*, 2018, 2019).

Neddylation of Cullins can provide a regulatory mechanism that fine-tunes substrate ubiquitination and targeting for proteasomal degradation. We found that Cand2 regulates protein levels of Grk5 predominantly independent of the transcript level. Previous work showed that following pressure overload, Grk5 accumulated in the nucleus of cardiomyocytes and acts in the nucleus as a class II histone deacetylase (HDAC) kinase, phosphorylating specifically HDAC5 leading to its nuclear export and de-repression of the transcription factor MEF2 as well as NFAT (Cannavo *et al*, 2016; Traynham *et al*, 2016). Furthermore, transgenic mice with Grk5 cardiac overexpression could not cope with pressure overload (Martini *et al*, 2008). Once free of repression, MEF2 and NFAT are responsible for the transcription of hypertrophic genes which leads to pathological hypertrophy (Zhang *et al*, 2011).

Here we found that reduced levels of Grk5 in Cand2 depleted myocytes resulted in decreased MEF2 activity. We also confirmed the direct interaction of Cand2 with Cullin1 and Grk5 in myocytes, supporting the hypothesis that Cand2 controls Cullin1-dependent Gr5k levels in cardiomyocytes. Moreover, pharmacological inhibition of neddylation using a selective neddylation inhibitor MLN4924 resulted in loss of Cullin1 neddylation and increased cell size, which was associated with increased Grk5 levels independent from transcript levels. Cardiomyocytes depleted of Cand2 did not respond to MLN4924, suggesting that Cand2 is necessary for growth induction caused by MLN4924. Rescue of Grk5 expression in Cand2 depleted cells reversed the observed phenotype with increased pathological cell size similar to control cells.

Taken together, we suggest a novel mechanism by which pathological mTORC1 signaling links translationally-controlled Cand2 expression to transcriptional activity (mRNA translation controls transcriptional activity). We suggest that these findings might be transferable toward novel treatment options. We hypothesize that Cand2 is a promising target as it is a cardiac muscle-specific protein, early (translationally) upregulated during disease initiation, and sufficient for pathological remodeling. Future studies will develop a nucleic acid-based therapy against Cand2. Recent advances in nucleic acid-based therapeutics show promising results in treatments for various diseases including pathological cardiac hypertrophy and it should be possible to target Cand2 translation by antisense oligonucleotides targeting the 5’ UTR of Cand2 (Morihara *et al*, 2017). Future studies are also needed to identify additional interaction partners of Cand2 dependent on the neddylation status of Cullin1 to understand how mTORC1-dependent upregulation of Cand2 affects the stability of protein networks in the nucleus.

## Material and Methods

The authors declare that all supporting data are available within the article (and in the Online Data Supplement).

Raw sequencing data have been made publicly available and can be accessed. RNA-Seq: Raw data have been uploaded to GEO (accession ID: GSE153364)

Ribo-Seq: Raw data have been uploaded to SRA (accession ID:SRP156230)

### Parallel Generation of Ribo-seq and RNA-seq Libraries

Ribo-seq and RNA-seq libraries were prepared for each biological replicate. Ribosome footprints were generated after immunoprecipitation of cardiac myocyte-specific monosomes with anti-HA magnetic beads after treating the lysate with RNase I. Libraries were generated according to the mammalian Ribo-seq kit (Illumina). Barcodes were used to perform multiplex sequencing and create sequencing pools containing at least eight different samples and always an equal amount of both RNA and RPF libraries. Sample pools were sequenced on the HiSeq 2500 platform using 50-bp sequencing chemistry.

The Ribo-Taq mouse model and ribosome-protected fragments immunoprecipitation were described previously (Doroudgar *et al*, 2019). To identify mTOR-dependent translatome *in vitro* polysome fractionations from NRCMs were isolated in a sucrose gradient according to standard procedure (Kmietczyk *et al*, 2019). The absorbance was monitored at 254 nm to record the polysome profile. The collected fractions were treated with RNaseI and mixed with Qiazol to recover ribosomal footprints. Generation of Ribo- and RNA-seq libraries and deepsequencing data processing were described in detail(Doroudgar *et al*, 2019). Briefly, 15 Million NRCMs were lysed in 500μl polysome buffer (20 mM Tris pH 7.4, 10 mM MgCl, 200 mM KCl, 2 mM DTT, 100 μg/ml CHX, 1% Triton X-100, 1U DNAse/μl) containing 100 μg/ml CHX. The lysate was used for RPF generation using polysome profiles after RNAse1 digestion. For complete lysis, the samples were kept on ice for 10 min and subsequently centrifuged at 20,000×g to precipitate cell debris and the supernatant was immediately used in the further steps. Sucrose solutions were prepared in polysome gradient buffer and 20 U/mL SUPERase-In (Ambion). Sucrose density gradients (10–50% wt/vol) were freshly made in SW40 ultracentrifuge tube using a BioComp Gradient Master (BioComp). Ribosome footprints were generated after treating the lysate with RNAse I (Ambion). Cell lysates were loaded onto sucrose gradients, followed by centrifugation for 250 min at 220,000×g, 4 °C, in an SW40 rotor. Separated samples were fractionated at 0.375 ml/min by using a fractionation system BioComp Gradient Station (BioComp) that continually monitors OD254 values. Monosomal fractions were collected into tubes at 0.3 mm intervals. Libraries were generated according to the mammalian Ribo-seq kit (Illumina). Sample pools were sequenced on the HiSeq 2000 platform using 50-bp sequencing chemistry.

### Data analysis and visualization

For RNA-Seq R and Bioconductor was used with the NGS analysis package systempipeR (Backman & Girke, 2016). Quality control of raw sequencing reads was performed using FastQC. Low-quality reads were removed using trim_galore (version 0.6.4). The resulting reads were aligned to mouse genome version mm10 from UCSC using tophat2 (Kim *et al*, 2013). EdgeR (version 3.26.8) was used to perform a differential expression analysis (Robinson *et al*, 2010). A false-positive rate of α= 0.05 with FDR correction was taken as the level of significance. Volcano plots were created using R (version 3.4.0) using the ggplot2 package (version 2.2.1).

For Ribo-seq data, only periodic fragment lengths were kept that showed a distinctive triplet periodicity. We used the automatic Bayesian selection of read lengths and ribosome P-site offsets (BPPS) method (Malone *et al*, 2016) to select and shift aligned reads to properly account for the P-site of the ribosome. We only consider data points with read count observations across all replicates. We used log2Fc >1.5 as a cutoff. For the myocytes data we do not estimate significance but provide only fold change values due to the absence of biological replicates.

GO-term enrichment analysis was performed using DAVID (Database for Annotation, Visualization and Integrated Discovery) with the rat genome as background. Only enriched GO terms with at least three significantly changed genes were kept for further analysis and ranked by Fisher Exact. Top enriched terms were retained and visualized with a custom plotting routine showing enrichment p-value.

### Mice, Surgery, and cardiac function

To study Cand2 *in vivo*, a novel Cand2 knock-out mouse line was created. An embryonic stem cell clone (EPD0169_1_A01-European Conditional Mouse Mutagenesis Program) was used to generate Cand2 knock-out mice. At 8 weeks of age, male Cand2^-/-^ mice underwent transverse aortic constriction (TAC; 27 gauge needle) or sham operation, as previously described (Kmietczyk *et al*, 2019;). Institutional Animal Care and Use Committee approval was obtained for all animal studies.

### Isolation and primary culture of neonatal and adult ventricular cardiomyocytes and neonatal mouse cardiomyocytes

Ventricular cardiomyocytes from 1 to 2 day old Wistar rats neonatal hearts (NRCMs) were prepared by trypsin digestion and percoll gradient separation according to standard procedures (Sanlialp *et al*, 2020). For analysis of hypertrophy, cells were treated with 50 uM PE for indicated time points. mTORC1 was pharmacologically inhibited with 250 nM Torin1 for 24h. For neddylation inhibition NRCMs were treated with 100 nM MLN4924 for 24h. Neonatal mouse heart myocytes were isolated from 1-day old wild-type and Cand2 knock-out mice by DNaseI/collagenase type III digestion according to the published protocol (Ehler *et al*, 2013). Adult ventricular cardiomyocytes were isolated using standard procedures as previously described (Kmietczyk *et al*, 2019).

### Plasmids and overexpressions

5’TOP-reporter constructs were obtained by cloning of Cand2, eEF2, and -Actin 5’UTRs into piCheck2 plasmid upstream of Renilla luciferase. HEK cells were transfected with 5’UTR-reporters for 24h using ViaFect reagent according to the manufacturer’s protocol (Promega). Next, HEK cells were treated with 250 nM Torin1 for 6h. Cand2 overexpression *in vitro* was obtained by NRCMs transduction with AAV6 vector. Mouse Cand2 (NM_025958.2) containing Myc-tag inserted after START codon was chemically synthesized (BioCat) and cloned into recipient vector pSSV9. To determine the optimal MOI, a preliminary dose-escalation experiment was conducted with different doses of Cand2 AAV6. For MEF2 and Grk5 studies, NRCMs were infected with adenoviruses harboring 3xMEF-luc (Firefly luciferase; a gift from Prof. Johannes Backs) and bovine Grk5 (gift from Prof. Philip Raake) for 24h.

### RNA interference

Pre-designed synthetic anti-rat Cand2 small interfering RNA (siRNA ID s140407), anti-Cul1 siRNA (s168977), anti-Rptor siRNA (s143005) and scrambled siRNA (Silencer™ Select Negative Control No. 1 siRNA) as negative control were purchased in Thermo Fisher Scientific. NRCMs were transfected with 25 nM final concentration of siRNAs by using HiPerfect transfection reagent according to the manufacturer’s instructions (QIAGEN).

### RNA isolation and RT-qPCR

Total RNA from NRCMs was isolated with Quick-RNA™ MiniPrep (Zymo Research) according to the manufacturer’s protocol. Liquid nitrogen snap-frozen tissues were homogenized in Precellys 24 homogenizer (Bertin Instruments) in 500 ul of lysis buffer containing 20 mM Hepes pH 7.4-7.5, 10 mM MgCl_2_, 200 mM KCl, 1% Triton, 1x protease inhibitor, 1x phosphatase inhibitor, 25 U/μl DNase I and 40U of RNasin. A 100ul aliquot was mixed with 1ml of Qiazol to isolate total RNA according to the standard protocols with chloroform and isopropanol precipitation. The quality of total RNA was checked on a NanoChip 2100 Bioanalyzer (Agilent). 100-500 ng of total RNA was reverse-transcribed into complementary DNA (cDNA) by using iScript™ Reverse Transcription Supermix (Biorad). Quantitative real-time PCR was performed using iTAQ™ SYBR Green PCR Kit (Biorad) according to the manufacturer’s instructions. Primers used in the study are shown in supplementary Table 1. Analysis of the specificity of the amplification product was performed by melting curve analysis. We calculated quantitative differences using the ΔΔC(T) method.

### Immunoblots

Samples were combined with the appropriately concentrated form of Laemmli sample buffer and then boiled before SDS-PAGE followed by transfer to PVDF membranes.

A list of all used antibodies is provided in Table 2.

### Histology, immunohistochemistry, and PLA

Sections were cut and deparaffinized using standard procedures. In brief, hearts were excised and embedded in formalin for 24 hours at room temperature. After cutting, sections were deparaffinized, rehydrated and antigens were *retrieved.*

NRCMs and ARCMs were plated on permanox or glass chamber slides (gelatin coated) and fixed by paraformaldehyde, permeabilized and blocked as described (Völkers *et al*, 2013). Cand2 primary antibody (Bethyl Laboratories) diluted 1:100 vol/vol in the respective blocking solution was applied to both types of slides overnight at 4°C. Cand2 was detected by FITC or Cy3-conjugated secondary antibody (Jackson Laboratories). Alexa Fluor™ 555 Phalloidin (Thermo Fisher Scientific) or conjugated phalloidin 633 (Jackson Laboratories) was used to detect F-actin. DAPI (Life Technologies) was diluted in Vectashield (Vectra Labs) mounting media and used as nuclear staining.

Proximity ligation assay was performed according to the manufacturer’s instructions (DuoLink in situ red kit Mouse/Rabbit) in rat and mouse neonatal cardiomyocytes. Primary antibodies were diluted 1:20 in DuoLink antibody diluent and incubated with cells at 4°C overnight. For Cand2-Grk5 detection anti-rabbit TIP120B and anti-mouse Grk5 were used. For Cand2-Cul1 anti-mouse TIP120B and anti-rabbit Cul1 antibodies were applied. The connective tissue was visualized with WGA Alexa Fluor™ 488 conjugate (Thermo Fisher Scientific). Images were obtained with 20x and 60x objectives.

### Immunoprecipitation

NRCMs were treated with 1uM MLN 4924 for 6h and lysed with ice-cold IP lysis buffer containing 40 mM Hepes pH 7.4, 2 mM EDTA, 10 mM sodium pyrophosphate, 10 mM glycerophosphate, and 0.3% CHAPS. M280 sheep anti-rabbit Dynabeads were coated with anti-rabbit TIP120B antibody in IP lysis buffer for 1h at room temperature. 500 ug of protein lysate was combined with antibody-coated beads and rotated at 4°C overnight. Beads were washed three times with IP lysis buffer supplemented with 150 mM NaCl. Immunocomplexes were eluted with SDS sample buffer by boiling for 5 min, and proteins were detected by immunoblotting with a corresponding antibody.

### Subcellular fractionation

Subcellular fractionation of C2C12 cells and left ventricles was performed as previously described with small modifications (Wysocka *et al*, 2001). Briefly, cells and tissues were washed with ice-cold PBS and homogenized in buffer A (10 mM Hepes pH 7.9, 10 mM KCl, 1.5 mM MgCl_2_, 340 mM sucrose, 10% glycerol, 1 mM DTT, 1x proteases inhibitor cocktails, 1X Triton) with tight-fitting disposable tissue grinder pestle and needled with 20 and 22 gauge needles. Nuclear pellets and cytoplasmic supernatants were separated by low-speed centrifugation at 1 300 x g, 4°C for 5 min. The final cytoplasmic fraction was pre-cleared by high-speed centrifugation at 20 000 x g for 5 min at 4°C. The nuclear pellet from low-speed centrifugation was dissolved in the hypotonic buffer B containing 3 mM EDTA, 0.2 mM EGTA, 1 mM DTT, 1x proteases inhibitor cocktails.

### Luciferase Assays

Cells were harvested 24–48 h after transfection/infection in passive lysis buffer (Promega). The samples were prepared according to the manufacturer’s protocol (Promega) and measured by using a LUMIstar OPTIMA plate reader. Luciferase’s signals were normalized to total protein amount determined with *RC-DC* kit (BioRad).

**Table 1.**
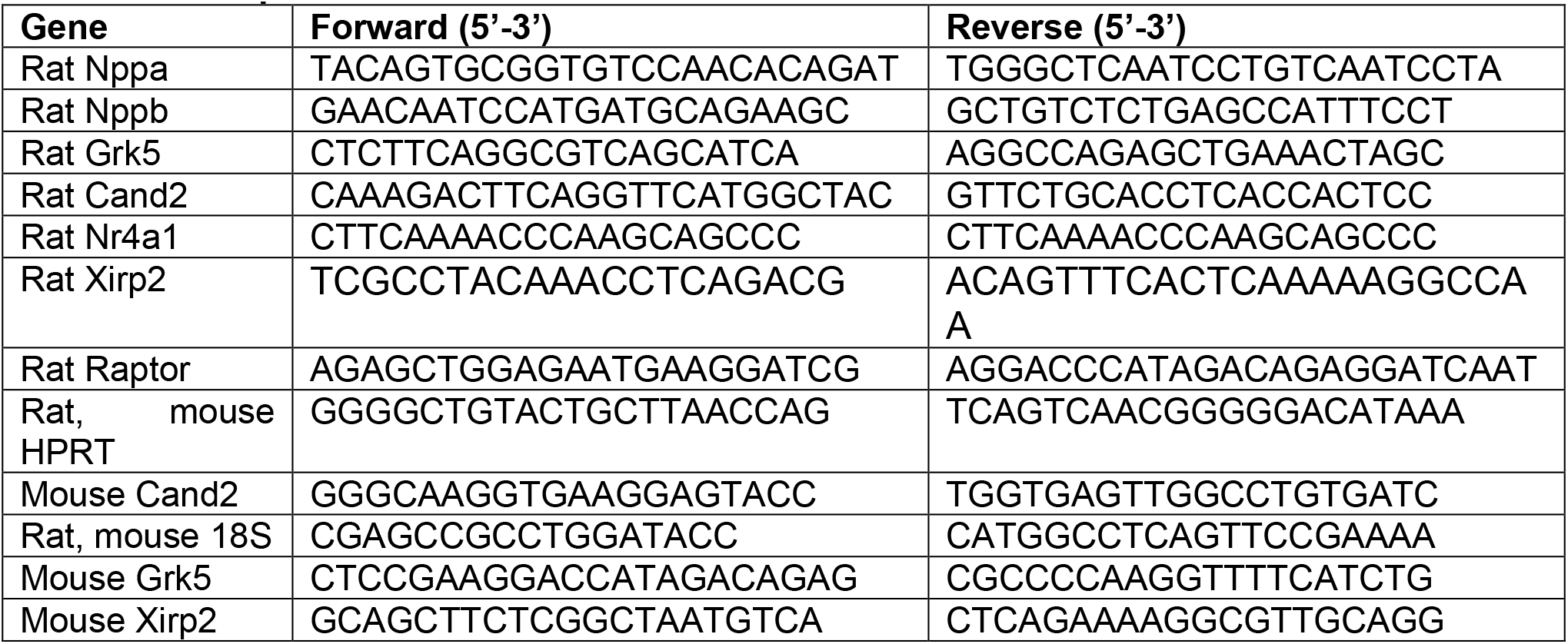
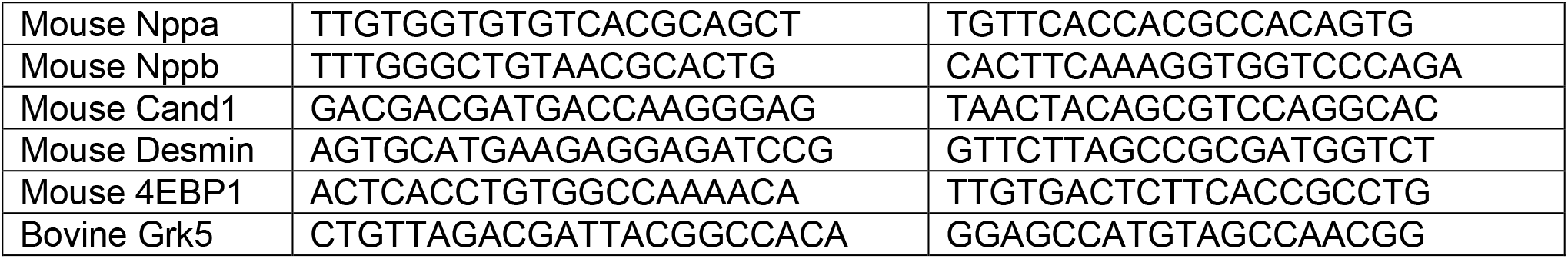
List of primers

**Table 2.** List of primary antibodies

| Antigen | Primary antibody | Origin | Company | Cat.# | Application |
| --- | --- | --- | --- | --- | --- |
| Cand2 | TIP120B | Rabbit | Bethyl Laboratories | A304-046A-M | WB, IP, IF, PLA |
| Cand2 | TIP120B (H7) | Mouse | Santa Cruz | sc-515406 | WB, PLA |
| Grk5 | Grk5 (D9) | Mouse | Santa Cruz | sc-518005 | WB, PLA |
| Cul1 | Cul1 (D5) | Mouse | Santa Cruz | sc-17775 | WB, PLA |
| Cul1 | Cul1 | Rabbit | Bethyl Laboratories | A303-373A-M | WB, PLA |
| GAPDH | GAPDH (G9) | Mouse | Santa Cruz | sc-365062 | WB |
| β-Actin | Beta-Actin (C4) | Mouse | Santa Cruz | sc-47778 | WB |
| pRib6S | pS6RP (Ser235/236) | Rabbit | Cell Signaling | 4858S | WB |
| p4E-BP1 | p4E-BP1 (Ser65) | Rabbit | Cell Signaling | 9451S | WB |
| 4E-BP | 4E-BP | Rabbit | Cell Signaling | 9452 | WB |
| pERK 1/2 | Thr218/Tyr220 | Rabbit | Cell Signaling | 3371 | WB |
| Myc-Tag | (71D10) | Rabbit | Cell Signaling | 2278S | WB |
| pAkt1 | Ser473 | Rabbit | Cell Signaling | 9271 | WB |
| Csq | calsequestrin | Rabbit | abcam | Ab3516 | WB |
| rpS5 | rpS5 | Rabbit | abcam | Ab210745 | WB |
| Lamin B1 | Lamin B1 (B10) | Mouse | Santa Cruz | sc-374015 | WB |
| Histone1 H3D | Histone1 H3D | Mouse | Santa Cruz | sc-134355 | WB |
| Cul4 | Cul4 (H11) | Mouse | Santa Cruz | sc-377188 | WB |
| Des | Desmin | Rabbit | Abcam | ab15200 | WB |
| eEF2 | eEF2 | Rabbit | Cell Signaling | 2332s | WB |
| rpS20 | rpS20 | Rabbit | Abcam | ab133776 | WB |

### Statistics

*In vivo* experiments were performed on 3-20 biological replicates (mice) for each treatment. Throughout the studies, the investigators were blinded to the sample group allocation during the experiment and analysis of the experimental outcome. Statistical analysis was performed using GraphPad Prism 7.0 (Graphpad Software Inc; www.graphpad.com) or R. All the data sets were tested for normality of distribution using the Shapiro-Wilks test (threshold *P*<0.05). For normally distributed data, values shown are mean±SEM. Statistical analysis of data involving two groups was performed using unpaired two-tailed t-test, for more than two groups 1-way ANOVA with the Bonferroni test applied to correct for multiple comparisons.Sequencing count data were modeled using negative-binomial distribution. For not normally distributed data a nonparametric test was used to test for significance between different groups. A Mann-Whitney test was performed when comparing two groups. A Kruskal-Wallis test was used when comparing multiple groups (more than two) followed by a Dunn’s multiple test comparison.

## Data Availability Section

Raw sequencing data have been made publicly available and can be accessed.

RNA-Seq: Raw data have been uploaded to GEO (accession ID: GSE153364) Ribo-Seq: Raw data have been uploaded to SRA (accession ID:SRP156230)

## Acknowledgements

A.A.G., H.A.K., M.V., S.D., and C.D. acknowledge the DZHK (German Center for Cardiovascular Research) Partner Site Heidelberg/Mannheim. C.D. acknowledges funding from the Klaus-Tschira Stiftung GmbH. S.D. acknowledges the European Society of Cardiology Basic Research Fellowship and the DZHK Excellence Programme. M.V. acknowledges the DFG (German Research Foundation, DFG VO 1659 2/1, DFG VO 1659 2/2, DFG VO 1659 4/1, DFG VO 1659 6/1), the Boehringer Ingelheim Foundation (Plus 3 Programme).

## Non-standard Abbreviations and acronyms

Cand2: Cullin-associated NEDD8-dissociated protein 2
CPM: Count Per Million
RPKM: Reads Per Kilobase Million
DEGs: Differential Expressed Genes
IP: Immunoprecipitation
mTOR: Mechanistic Target Of Rapamycin
nt: Nucleotide
NRCMs: Neonatal Rat Cardiac Myocytes
LV: Left ventricle
RPF: Ribosome Protected mRNA Fragments
Ribo-seq: Ribosomal sequencing
RNA-seq: RNA sequencing
TOP-motif: Terminal oligo pyrimidin motif
5’UTR: 5’ Untranslated Region
IRES: Internal Ribosomal Entry Site
TAC: Transverse Aortic Constriction
PE: Phenylephrine
Cx: Cycloheximide
PLA: Proximity Ligation Assay
WT: Wild-type
KO: Knock-out
KD: Knock-down
OE: Overexpression

## Expanded View Figures

**EV 1. Specific inhibition of mTOR pathway by Torin1.**

**(A)** NRCMs treatm with increasing concentration of Torin1 for 24h analyzed by immunoblotting. mTORC1 inhibition was detected by total and phosphorylated levels of S6RP and 4E-BP1. mTOR2 activity was analyzed by Akt phosphorylation. PI3K pathway was analyzed by the phosphorylation level of its downstream target Erk1/2. **(B)** Expression of mTOR-dependent transcripts. eEF2 and Rps5 levels were analyzed by immunoblot (left panel) and RT-qPCR (right panel) after PE and Torin1 treatment. Csq - calsequestrin has been used as a housekeeping protein. **(C)** mTOR-dependent expression of hypertrophy markers. RT-qPCR analysis of *Nppa* and *Nppb* mRNA levels after PE and Torin 1 treatment of NRCMs. **(D)** Polysome profiles of control and PE- and Torn1-treated NRCMs.

**EV 2**

**(A)** Representative bright-field images of isolated cardiac myocytes stimulated with (PE, 50 μM for 24 h) with or without mTOR inhibition (Torin 1,150 nM) and quantification of cell surface area measurements from control, PE treatment, and Torin 1 treatment. n=4 independent experiments, *-*P* < 0.01 different from PE. Scale bar 100μm **(B)** Luciferase activity from control, PE treated, and Torin 1 treated cardiac myocytes. n=2 independent experiments, *-*P* < 0.01 different from PE. **(C)** Heart weight (HW) to body weight (BW) ratio (HW/BW) in control and Torin 1-treated mice (10 mg/kg BW) 2 days after sham or TAC surgery. *-*P* < 0.05 different from TAC. n=3 per group

**EV 3. Cand2 knock-out mouse model characterization.**

**(A)** Cand2 protein levels in WT, heterozygous, and homozygous Cand2 knock-out mice analyzed by immunoblotting in muscle tissues. **(B)** The percentage of ejection fraction in WT and Cand2 KO analyzed by echocardiography prior sham/TAC surgeries. **(C)** The baseline of LV weight (mg) to body weight (g) ratios of WT and KO Cand2 mice. **(D)** RT-qPCR analysis of Cand2 and Cand1 transcript levels in WT and Cand2 KO mice. **(E)** *Nppa* and *Nppb* **(F)** transcripts in WT and Cand2 KO mice.

**EV 4. Cand2 interacts with Cul1 and Grk5 in cardiomyocytes.**

**(A)** Cand2, Cul1, and Grk5 association analyzed by co-immunoprecipitation (IP) of endogenous Grk5 with Cand2-specific antibody and immunoblotting (i – input; IgG – negative control, IP with anti-rabbit IgG antibody) and Immunoprecipitation of endogenous Cul4 with Cand2 specific antibody (i – input; IgG – negative control, IP with anti-rabbit IgG antibody). **(B)** Interaction of Cand2 with Grk5 measured by proximity ligation assay (PLA). PLA (red) in primary neonatal rat (NRCMs) and cardiomyocytes isolated from hearts of wild-type **(C)** and Cand2 KO **(D)** neonatal mice (NMCMs). Each red dot represents the interaction of endogenous Cand2 and Grk5. No dots were detected in mouse cardiomyocytes from Cand2 KO hearts. **(E)** PLA of Cand2 and Cullin1 in NRCMs and **(F)** NMCMs from WT and **(G)** Cand2 KO mice. Red dots represent the Cand2 and Cul1 association. No dots were detected in mouse cardiomyocytes from Cand2 KO hearts. Scale bar 20μm

**EV 5. Cand2 regulates Grk5 protein level by Cullin1 neddylation inhibition.**

**(A)** RT-qPCR analysis of Cand2 mRNA levels in NRCMs after neddylation inhibition (MLN) and Cand2 KD with siRNA. **(B)** Cul1 expression in NRCMs overexpressing (left graph) and depleted of Cand2 (right graph) examined by RT-qPCR. **(C)** Grk5 and Cul1 mRNA levels in NRCMs overexpressing Grk5 measured by RT-qPCR.

